# Higher-Order Mechanical Neighbourhood Sensing Governs Cell Fate in Complex Tissues

**DOI:** 10.64898/2026.08.13.744616

**Authors:** Alphy John, Tianxiang Ma, Duncan Mackay, Elizabeth Catherine Connolly, Julien Colombani, Amin Doostmohammadi, Ditte S. Andersen

## Abstract

Adult tissues maintain precise architectures composed of multiple interspersed cell types, yet prevailing models of cell fate patterning rely on pairwise interactions between neighbouring cells that generate binary decisions. How local interactions encode higher-order, beyond pairwise, multicellular organization remains unclear. Here, using the *Drosophila* midgut as a model of adult tissue self-organization, we identify a higher-order neighbourhood-sensing mechanism that couples cell identity, mechanics and stem cell (SC) fate. We show that the receptors Cirl/Latrophilin and Toll-8 are expressed in complementary cellular compartments, restricting their engagement to heterotypic interfaces between progenitors and differentiated enterocytes (ECs). These interactions generate interface-specific myosin II-dependent tension that depends on EC neighbourhood composition and mechanically gates Delta–Notch signalling. Thus, although Delta–Notch acts through pairwise interactions, these fate decisions are modulated by a higher-order mechanical state integrating information from surrounding cell contacts. A mathematical model incorporating this coupling reproduces multicellular tissue architecture from first principles and predicts that loss of higher-order coupling destabilizes progenitor organization. This prediction is consistent with the excessive fate transitions, topological disorganization and regenerative defects observed after Cirl–Toll-8 disruption. These findings establish higher-order mechanical neighbourhood sensing as a general principle by which tissues integrate local cellular identities to maintain and restore complex architectures.

## Introduction

Adult tissues maintain precise architectures composed of multiple interspersed cell types. Achieving this organization depends on assembling the right composition of cells but also precise coordination of their positioning. How cells use local information to establish and preserve these arrangements, and to restore them after injury, remains a fundamental problem in biology.

Delta-Notch mediated lateral inhibition provides a canonical framework for local cell fate patterning. Through pairwise signalling, Delta-expressing cells activate Notch signalling in neighbouring cells, which represses Delta expression and drives adjacent cells toward opposing fates, producing stable salt-and-pepper patterns (Collier, Monk et al. 1996, Bray 2006, Kovall, Gebelein et al. 2017, Henrique and Schweisguth 2019, Sprinzak and Blacklow 2021). While this framework captures binary fate choices (Shaya and Sprinzak 2011, Sjoqvist and Andersson 2019, Bocci, Onuchic et al. 2020), it does not address how these decisions are influenced by additional cell types within complex tissue environments. Such context could give rise to higher-order interactions, where the outcome of a pairwise fate decision is modified by the composition of the surrounding neighbourhood. Whether tissues use higher- order interactions to organize multicellular architecture, and how they are physically implemented, remain unknown.

Addressing how higher-order neighbourhood interactions are physically implemented requires a tissue in which spatial organisation of multiple cell types is stereotyped, quantifively accessible, and can be tracked through cycles of disruption and repair. Adult tissues with high regenerative capacity enable direct interrogation of collective cell behaviour, as injury induces transient loss of spatial organization that must be reconstituted to restore homeostasis. Flat epithelia are particularly well suited to this analysis because collective dynamics emerge primarily from direct cell–cell interactions rather than signalling gradients and spatial constraints characteristic of structured niches (Mesa, Kawaguchi et al. 2018). In tissues, such as the respiratory epithelium and interfollicular epidermis, uniform SC distribution across the tissue ensures an even production of differentiated progeny (Blanpain and Fuchs 2009, Tata and Rajagopal 2017, Hewitt and Lloyd 2021). Here we use the Drosophila midgut, a flat monolayer epithelium with a stereotyped spatial cellular pattern, to determine how cells sense their neighbourhood composition, how this information shapes fate decisions, and whether it is required to restore tissue architecture after injury.

The adult Drosophila gut provides a well-defined system for investigating how spatial patterns are established and maintained. Under homeostatic conditions, intestinal SCs (ISCs) and their immediate progeny, enteroblasts (EBs), form pairs that are evenly spaced throughout the epithelium (Micchelli and Perrimon 2006, Ohlstein and Spradling 2007, Bardin, Perdigoto et al. 2010, de Navascues, Perdigoto et al. 2012). ISC-EB pairs arise when ISCs divide asymmetrically giving rise to one Delta-positive ISC and one Notch positive EB, which can subsequently mature into an EC (Micchelli and Perrimon 2006, Ohlstein and Spradling 2006, Ohlstein and Spradling 2007, Guo and Ohlstein 2015, Guo, Huang et al. 2019). While Delta-Notch lateral inhibition establishes binary ISC-EB fate decisions, it cannot account for how these pairs achieve and maintain their stereotyped spacing within a multicellular environment, leaving unresolved the mechanisms that couple local fate decisions to higher-order spatial organization.

Here we identify the evolutionary conserved adhesion G protein-coupled receptor (aGPCR), Cirl/Latrophilin and the Toll receptor, Toll-8, as critical regulators of higher-order neigbourhood sensing, shaping adult tissue architecture. In the Drosophila intestinal epithelium, their complementary expression enables progenitors to sense the identity of surrounding enterocytes and couple neighbourhood composition to Delta-Notch dependent fate decisions through cortical tension, implementing a higher-order interaction in which the outcome of a pairwise fate is set by the surrounding cellular neighbourhood. Genetic perturbations and mathematical modelling show that this coupling stabilizes progenitor organization and is required to restore spatial order after injury. These findings demonstrate how Cirl–Toll-8 interactions translate differences in cell identity into mechanical information, ensuring both steady-state tissue organization and the faithful re-emergence of spatial pattern after injury.

## Results

### Complementary Cirl and Toll-8 expression encodes spatial identity

The Drosophila midgut exhibits a stereotyped spatial architecture in which ISC-EB pairs are evenly distributed throughout the epithelium, each separated by one to two mature enterocytes (ECs) (Fig. 1a-d). Maintaining this arrangement requires progenitors to distinguish contacts with ECs from contacts with other progenitors. Because long Toll receptors mediate interface recognition during embryogenesis (Pare, Vichas et al. 2014, Pare and Zallen 2020), we asked whether they perform a similar function in the adult intestine.

**Figure 1:**
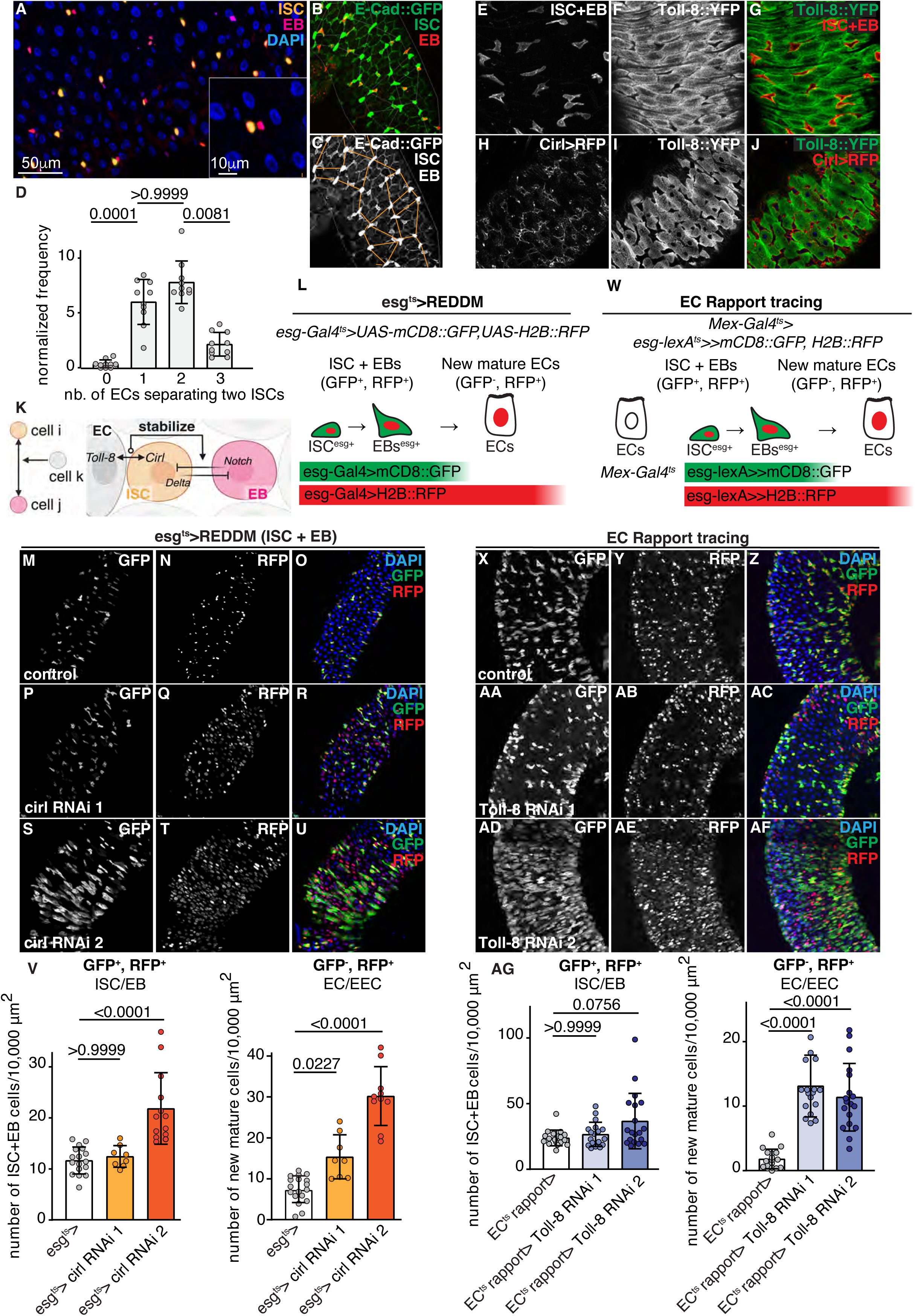
Complementary Cirl and Toll-8 expression encodes spatial identity. (a-d) Confocal images showing the stereotypic distribution of ISC-EB pairs in the posterior midgut. ISCs are typically separated by 1-2 ECs as quantified by “EC hop-distance” analyses in d. (e-g) Confocal images of posterior midguts from flies expressing *escargot (esg*)>RFP to label ISCs and EBs in red and Toll-8::YFP from its endogenous promoter. (h-j) Confocal images of posterior midguts from flies expressing *cirl>UAS-RFP* together with endogenous Toll-8::YFP reveal complementary expression patterns, with Cirl restricted to ISCs and EBs and Toll-8 to ECs. (k) Model for how Cirl–Toll-8 interactions at progenitor–EC interfaces enable higher-order neighbourhood sensing to coordinate ISC–EB pairs and stabilize their stereotyped spatial organization. (l) Schematics of the ReDDM system. Membrane-tethered CD8::GFP and nuclear H2B::RFP are co-expressed in ISCs and EBs under the *esg>* driver. Upon differentiation, esg-Gal4 activity is lost, terminating reporter expression; GFP is rapidly degraded whereas RFP persists in newly differentiated progeny. (m-v) ISC-EB-specific knockdown of Cirl increases tissue turnover. Representative confocal images of posterior midguts from control (m-o) and ISC-EB-specific Cirl knockdown (p-u) flies are shown. In all ReDDM experiments, RFP⁺/GFP⁻ polyploid cells were scored as newly generated ECs (v, n = 18, 8, 13). (w) Schematic of the EC-Repressible activity paracrine reporter (Rapport) tracing system. The UAS-Gal4 system is used to deplete Toll-8 in ECs, while tissue turnover is monitored through LexA/Aop-based lineage tracing. Membrane-tethered CD8::GFP and nuclear H2B::mCherry::HA are co-expressed in ISCs and EBs under the *esg>lexA* driver. Upon differentiation, *esg>lexA* activity is lost, terminating reporter expression; GFP is rapidly degraded whereas RFP persists in differentiated progeny. (x-af) Representative confocal images of posterior midguts from control and EC-specific Toll-8 knockdown flies are shown. In all EC-Rapport experiments, RFP⁺/GFP⁻ polyploid cells were scored as newly generated ECs (ag; n = 18, 18, 19). Statistical tests: Kruskal–Wallis with post-hoc multiple comparison analysis. Data are presented as mean values ± SD.

We found that Toll-8, but not Toll-2, Toll-6, or Toll-7, is expressed in the posterior midgut epithelium (Fig. 1f-g, i-j and S1a-f). Strikingly, Toll-8 expression is confined to ECs and excluded from stem and progenitor cells (Fig. 1e-g and S1g). Conversely, Cirl/Latrophilin, previously identified as a binding partner of Toll-8 on neighbouring cells (Lavalou, Mao et al. 2021, Scholz, Dahse et al. 2023), exhibited a complementary expression pattern, with high expression in ISCs and EBs and little or no expression in ECs (Fig. 1h,j). This mutually exclusive expression pattern means that every progenitor-to-EC boundary is marked by heterotypic Cirl-Toll-8 contact, while boundaries between two progenitor cells carry no such signal. We hypothesized that Cirl and Toll-8 provide a molecular code through which progenitors can distinguish enterocyte contacts from contacts with other progenitors and adjust tissue turnover accordingly (Fig. 1k).

To test this, we used the ReDDM and Rapport tracing systems (Antonello, Reiff et al. 2015, Zipper, Corominas-Murtra et al. 2025) to knock down Cirl in ISCs and EBs and Toll-8 in ECs, respectively, which caused an increase in tissue turnover in the absence of damage (Fig. 1l–ag). Moreover, ISC-specific Cirl depletion expanded the EBs pool without altering ISC cell numbers (Fig. 2a–k), indicating that loss of heterotypic receptor engagement promotes asymmetric ISC-EB divisions and accelerates EB fate acquisition. Thus, complementary Cirl and Toll-8 expression not only distinguishes heterotypic interfaces but also restrains progenitor fate transitions during homeostasis.

**Figure 2:**
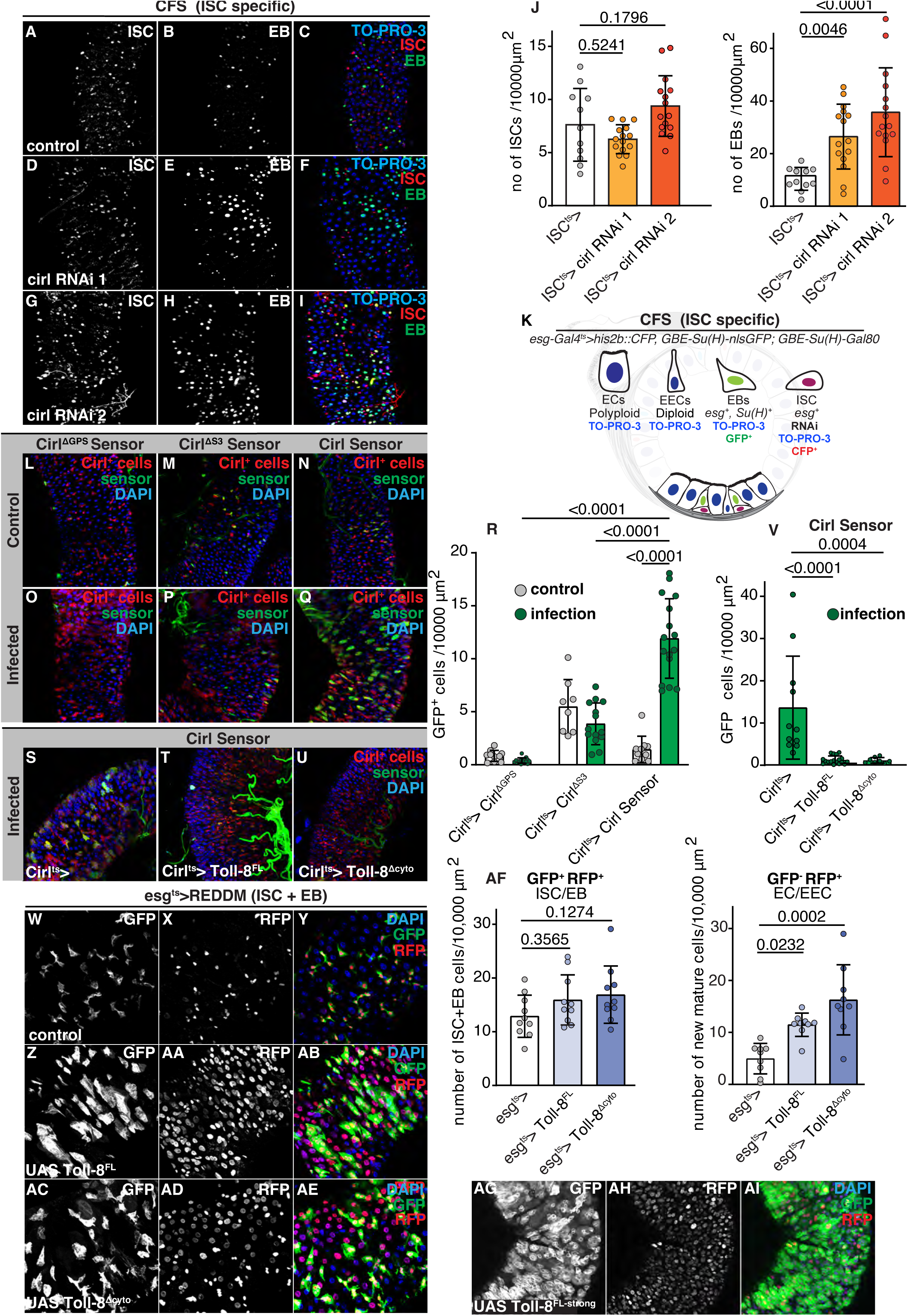
Trans-activation of Cirl by Toll-8 requires spatial segregation. (a-i) Representative confocal images of posterior midguts from control (a-c) and ISC-specific Cirl knockdown (d-l) flies are shown. ISC-specific Cirl depletion increases the number of EBs, while maintaining an unchanged ISC population as quantified in (j, n = 11, 15, 15). (k) Schematic of the Cell Fate Sensor, CFS^ISCts^ system used to distinguish midgut cell populations. Stem cells are visualized by expressing nuclear localized UAS-His2b::CFP by combining Su(H)GBE:Gal80 with esg-Gal4, EBs are marked by nuclear Su(H) GBE:nlsGFP, while the nuclei of all cells are labelled by TO-PRO-3. (l-r) Global epithelial damage induced by a 16-hour oral Ecc15 infection increases Cirl activity. (l-q) Confocal images showing that guts expressing the Cirl activity reporter under the control of the endogenous *cirl* promoter exhibited increased reporter activity following oral *Pe* infection, as quantified in (r, n=10, 15, 8, 14, 10, 16, 11, 18, 8). This infection-induced increase was abolished in a cleavage-deficient reporter mutant that cannot undergo proteolytic processing (Cirl^ΔGPS^, Cirl^ΔS3^, o-p). (s-v) Similarly, ISC-EB-specific expression of full-length Toll-8 (Toll-8^FL^, t) or Toll-8 lacking the intracellular domain (Toll-8^Δcyto^, u) suppressed infection-induced Cirl activity as quantified in v. Confocal images of posterior midguts from control flies (w-y) or flies with ectopic ISC-EB-specific expression of Toll-8^FL^ (z-ab, ag-ai) or Toll-8^Δcyto^ (ac-ae). ISC-EB-specific expression of Toll-8^FL^ or Toll-8^Δcyto^ increases tissue turnover as quantified in af. In all ReDDM experiments, RFP⁺/GFP⁻ polyploid cells were scored as newly generated ECs (n = 10, 10, 10). Statistical tests: Kruskal–Wallis with post-hoc multiple comparison analysis. Data are presented as mean values ± SD.

### Trans-activation of Cirl by Toll-8 requires spatial segregation

Cirl belongs to the adhesion GPCR family, whose members are activated by mechanical force following autocleavage of their extracellular domain. Previous studies have shown that trans-interaction with Toll-8 promotes this activation, while cis-expression of both proteins in the same cell inhibits it (Lavalou, Mao et al. 2021, Scholz, Dahse et al. 2023). Because Cirl and Toll-8 are normally confined to opposing cellular compartments, we asked whether this spatial segregation is required for productive Cirl engagement.

Using a Cirl activity sensor under its endogenous promotor to track force-dependent Cirl activation (Cirl>UAS-Cirl Sensor; (Scholz, Dahse et al. 2023)), we found that Cirl activity was low and sporadic under homeostatic conditions but became widespread following infection-induced epithelial damage (Fig. 2l-r). This activation was strongly suppressed when full-length Toll-8 (Toll-8^FL^) or Toll-8 lacking the intracellular domain (Toll-8^Δcyto^) was mis-expressed in ISCs and EBs in cis (Fig. 2s-v), confirming that Toll-8 functions as a ligand for Cirl and that productive engagement is strictly trans-dependent. Accordingly, ISC-EB specific overexpression of Toll-8^FL^ or Toll-8 ^Δcyto^ increased tissue turnover, phenocopying Cirl loss of function (Fig. 2w-af). Strikingly, strong UAS-driven Toll-8 mis-expression in ISCs and EBs disrupted epithelial patterning entirely, producing large tumours (Fig. 2ag-ai; *esg>Toll8^FL-strong^*). Collectively, these findings show the spatial segregation of Cirl and Toll-8 to opposing compartments is required to restrict productive receptor engagement to progenitor-EC interfaces, and that subverting this segregation is sufficient to disorganize the tissue.

### Heterotypic receptor engagement generates interface-specific mechanical tension to scale Notch signalling to neighbourhood composition

During embryogenesis, Toll-8-Cirl trans-interactions promote Cirl-dependent activation of Myosin II (Myo-II) at Toll-8 domain boundaries, guiding planar actomyosin polarization for anterior-posterior convergent extension (Lavalou, Mao et al. 2021). Similarly, ISCs and EBs displayed markedly elevated cortical Myo-II relative to surrounding ECs, with enrichment concentrated at progenitor-to-EC interfaces (Fig. 3a-c, S2a-c). Depleting Cirl in ISCs or Toll-8 in ECs, significantly reduced this enrichment (Fig. 3d-n), suggesting that interface-specific tension depends on heterotypic receptor engagement across compartment boundaries. Loss of Cirl caused ISCs to acquire an enlarged, spread morphology characteristic of activated stem cells, supporting a model in which Cirl–Toll-8 contacts at progenitor–EC interfaces generate the local mechanical asymmetry that maintains progenitor quiescence (Fig. S2d-g).

**Figure 3:**
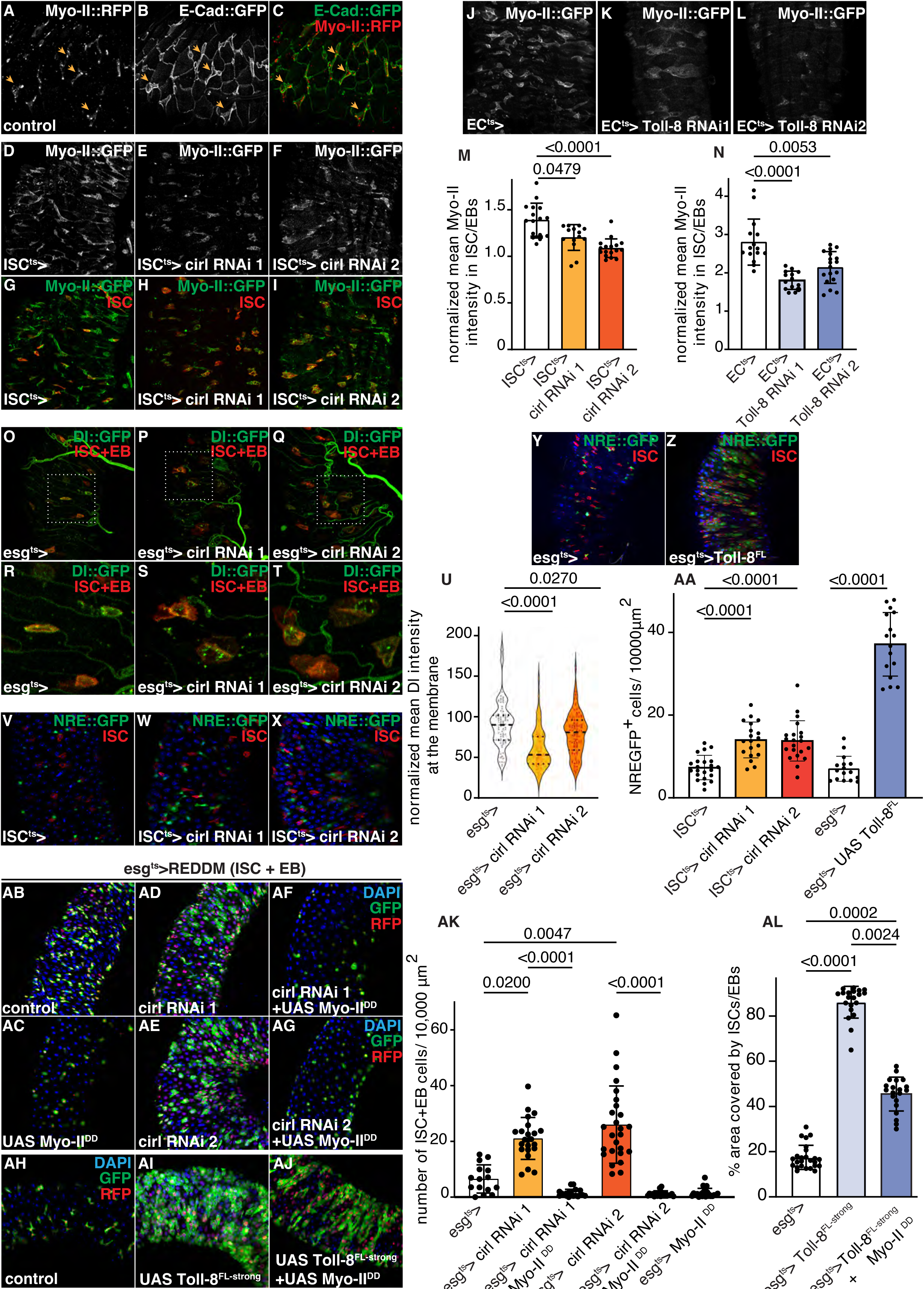
Cirl–Toll-8 signaling maintains cortical tension to restrain Notch activity. (a-l) Confocal images of posterior midguts dissected from flies expressing endogenous Myo-II::RFP and E-Cad::GFP show elevated cortical Myo-II at interfaces between ISC-EB pairs and neighboring ECs (a-c; yellow arrows). (d-l) Confocal images of posterior midguts dissected from flies expressing endogenous Myo-II::GFP combined with ISC^ts^>RFP (d-i) or *mex^ts^*> (j-l). ISC-specific (ISC^ts^>) Cirl depletion (d-i) or EC-specific (*mex^ts^>*) Toll-8 depletion (j-l) reduces cortical Myo-II levels as quantified in m-n (n = 16, 15, 17, 15, 14, 18). (o-t) Confocal images of posterior midguts expressing endogenous Delta::GFP. ISC-EB-specific Cirl depletion increases Delta internalization into intracellular vesicles resulting in reduced membrane-associated Delta as quantified in u (n = 84, 67, 94 cells). Posterior midguts from flies expressing the Notch activity reporter, NRE::GFP, diplay elevated Notch signaling following ISC-EB-specific Cirl depletion (v-x) or Toll-8-FL overexpression (y-z) or, as quantified in aa (n = 22, 19, 21, 16, 16). (ab-al) Increased cellular tension suppresses the tissue turnover induced by Cirl depletion. (ab-ag) Tissue turnover was assessed using the ReDDM system following ISC-EB-specific Cirl depletion, Myo-II^DD^ overexpression, or their combined expression. Co-expression of Myo-II^DD^ efficiently suppressed the increase in tissue turnover induced by Cirl knockdown (af, ag) or Toll-8^FL^ overexpression (ah-aj), as quantified in ak-al (n = 15, 22, 18, 26, 20, 24). Kruskal–Wallis with post-hoc multiple comparison analysis. Data are presented as mean values ± SD.

Tissue renewal in the absorptive lineage is governed by two sequential Notch thresholds: a low-threshold event that specifies EB fate in progenitor daughter cells, and a high-threshold event that drives terminal EC differentiation (Antonello, Reiff et al. 2015, Martin, Sanders et al. 2018). Endocytosis of Delta in signal-sending cells exerts a pulling force on Notch in receiving cells, triggering a proteolytic cleavage cascade that releases the Notch intracellular domain and drives transcriptional reprogramming (Seib and Klein 2021). We asked whether Cirl-Toll-8 generated tension suppresses Delta internalization and thereby controls the rate at which progenitors cross these thresholds. Reducing Cirl activity, either through ISC-EB specific Cirl depletion (Fig. 3o-u) or Toll-8^FL^ overexpression (S2h-m), increased Delta::GFP internalization and elevated Notch reporter activity (Fig. 3v-aa; NRE:GFP; (Saj, Arziman et al. 2010).

To establish causality between tension and Notch signalling independently of the receptor system, we manipulated Myo-II directly. Myo-II depletion phenocopied Cirl loss, enhancing Delta::GFP endocytosis and Notch activity (Fig. S2n-x). Notably, Myo-II activation (*esg>* Myo-II*^DD^*; (Mitonaka, Muramatsu et al. 2007)) was sufficient to rescue the elevated tissue turnover caused by Cirl depletion or Toll-8 overexpression (Fig. 3ab-al), placing Myo-II downstream of Cirl in the pathway restraining fate transitions. Collectively, these findings show that progenitors fine-tune their Notch activity levels through the mechanical output of Cirl-Toll-8 engagement: the greater the fraction of EC neighbours, the higher the cortical tension, and the lower the rate of fate conversion. The pairwise Notch outcome between two progenitor cells is thus modified by the composition of their surrounding neighbourhood.

### A mechanochemical model of higher-order cell interactions reproduces tissue architecture and defines a progenitor stability boundary

To test whether higher-order neighbourhood coupling is sufficient to account for tissue-level patterning, we developed a mechanochemical vertex model comprising ISC-like, EB-like, and EC-like cells (Fig. 4a). Vertex models represent tissue as a network of polygonal cells whose vertices move under mechanical forces derived from an energy functional (Guerrero, Perez-Carrasco et al. 2019, Bajpai, Chelakkot et al. 2022, Bocanegra-Moreno, Singh et al. 2023, Ma, Bonn et al. 2026). Progenitor-progenitor interactions follow conventional pairwise Delta-Notch lateral inhibition. The higher-order term enters through Cirl-Toll-8 mechanosensing: for each progenitor, the fraction of its contacts formed with ECs determines a local heterotypic-contact signal. This signal is multiplied by a mechanosensing gain (*g*) and converted into increased cortical contractility, reducing the effective Delta signal available for Notch activation. A progenitor surrounded by ECs therefore experiences weaker Notch signalling than one surrounded by other progenitors, meaning that pairwise fate decisions are modulated by the shared EC neighbourhood, implementing a higher-order interaction in the formal sense (see Methods for simulation details and Fig. S3a-b for sensitivity analyses).

**Figure 4:**
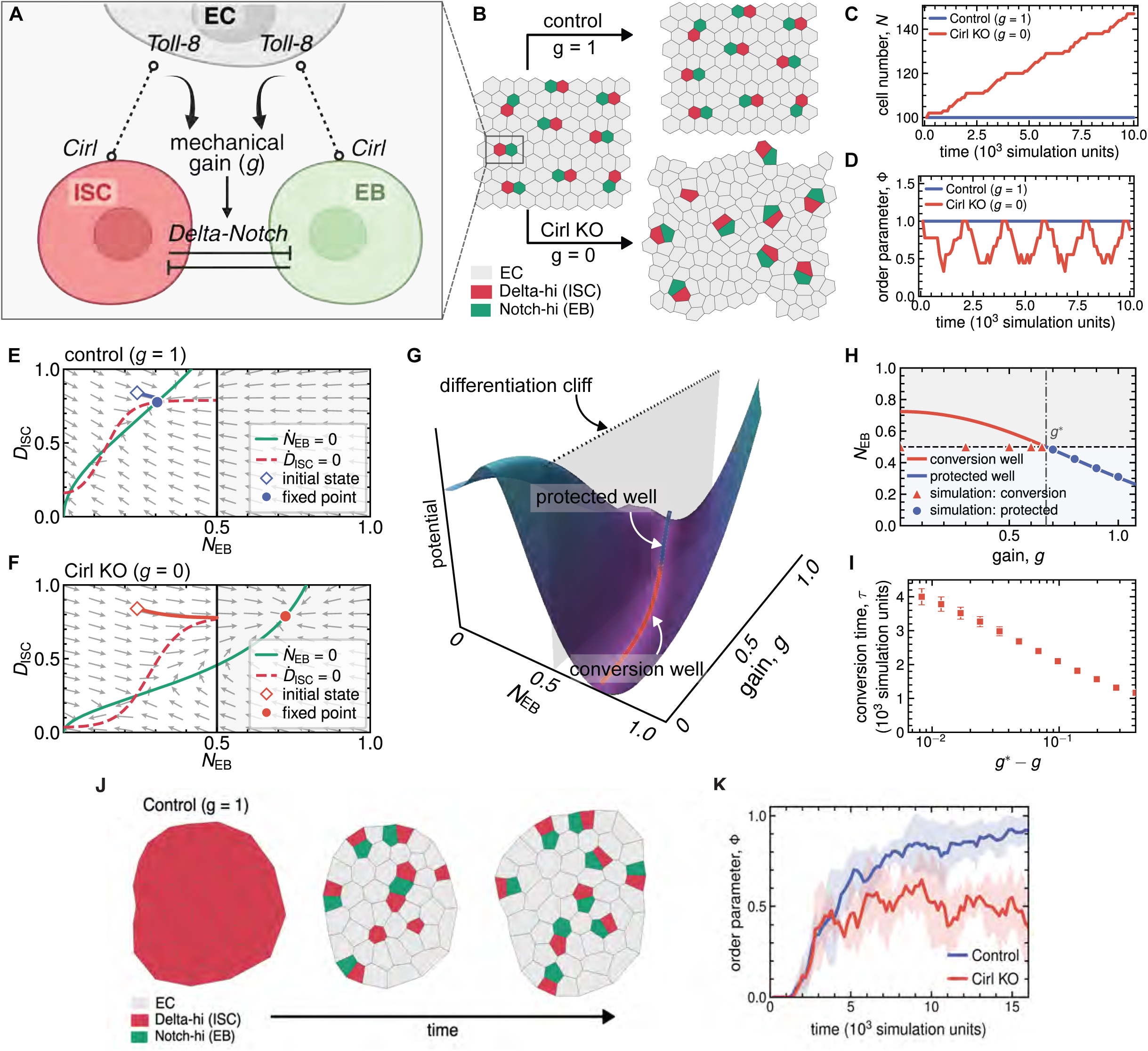
Cortical tension gates Delta-Notch signalling to stabilize cellular patterns. (a) Schematic of the model. Cirl–Toll-8 interactions at progenitor–EC interfaces generate a mechanical signal whose strength depends on the mechanosensing gain, g. This signal modulates cortical tension at the ISC–EB interface, making Delta–Notch coupling mechanosensitive and allowing the surrounding EC neighbourhood to influence progenitor fate. (b) Representative vertex model configurations at the initial state and after simulation with intact mechanosensing (g = 1, control) or complete loss of mechanosensing (g = 0, Cirl knockout). Control tissues maintain regularly distributed ISC–EB pairs; loss of coupling leads to progressive disruption of progenitor organization. ECs are shown in grey, Delta-high ISCs in red, and Notch-high EBs in green. (c, d) Time evolution of total cell number, N (c), and the progenitor-pattern order parameter, φ (d), for control and Cirl-knockout simulations. Intact coupling maintains both cell number and tissue organization; loss of coupling causes continued cell production and recurrent pattern disruption. (e, f) Reduced phase portraits for a single ISC–EB pair with intact coupling (g = 1; e) or no coupling (g = 0; f). Arrows show the local flow in Delta–Notch state space. Green and red curves denote the EB and ISC nullclines, respectively. Diamonds mark the initial state used in panel b; filled circles mark the stable fixed point. The vertical line marks the EB differentiation threshold. With intact mechanosensing, the fixed point stays below this threshold; loss of coupling shifts it into the differentiation regime. (g) Waddington-like potential landscape obtained by continuously varying g, with a protected progenitor well and a conversion well separated by a differentiation cliff. The orange trajectory traces the shift in the stable state as mechanosensing is reduced. (h) Steady-state EB activity as a function of g. A critical gain, g*, separates a protected progenitor state from a conversion state. Symbols show full-model simulations classified as protected or converting; solid curves give the corresponding reduced-model solutions. The horizontal dashed line marks the differentiation threshold. (i) Conversion time, τ, as a function of the distance from the critical point, g* − g. Conversion slows markedly as g approaches g*, a signature of critical slowing near the progenitor-stability boundary. Data are mean values across 3 independent simulations. (j) Representative simulations of tissue regeneration in the mechanochemical vertex model with intact Cirl–Toll-8 mechanosensing (control). Starting from a progenitor-enriched state, the tissue progressively regenerates a spatially dispersed pattern of Delta-high ISCs and Notch-high EBs surrounded by ECs. (j, k) Time evolution of the pattern order parameter, φ, in control and Cirl-knockout simulations. Intact mechanosensing promotes progressive recovery of tissue organization, whereas loss of Cirl reduces and destabilizes pattern restoration. Lines show mean values across 6 independent simulations.

Starting from a homeostatic configuration, the model maintained stable progenitor patterning when higher-order coupling was intact (*g* = 1) and reproduced progressive pattern loss when it was removed (*g* = 0), recapitulating the Cirl-depletion phenotype (Fig. 4b-d). Reducing the model to a single ISC-EB pair surrounded by EC neighbours revealed the mechanistic basis: higher-order coupling keeps Notch activity below the differentiation threshold, producing a stable fixed point that preserves the progenitor pair, while without coupling the fixed-point shifts beyond the threshold and the pair is lost (Fig. 4e-f). Continuously varying *g* constructed a Waddington-like stability landscape with a sharp boundary separating stable progenitor maintenance from progressive differentiation, predicting that small reductions in neighbourhood sensing near this boundary disproportionately destabilize progenitor identity (Fig. 4g-i). Higher-order neighbourhood coupling is therefore sufficient to account for stable multi-cell-type tissue architecture, and the stability boundary defined by the model constitutes a quantitative prediction for how sensitively tissue organization responds to partial loss of Cirl-Toll-8 signalling. The model further predicts a role for higher-order coupling in tissue recovery following spatial disruption. When initialized from a disrupted, progenitor-dense configuration lacking positional information, the model spontaneously restored the spatial pattern through an initial burst of asymmetric divisions followed by convergence towards the homeostatic arrangement. In contrast, removing higher-order coupling resulted in persistent progenitor clustering without recovery of spatial order (Fig. 4j-k; Movies 1-2). This predicts that Cirl depletion should specifically impair repatterning after injury while preserving the initial damage response.

### Higher-order neighbourhood sensing directs symmetric stem cell divisions and tissue repatterning after injury

We next asked whether higher-order coupling also governs the tissue’s response to injury. Global epithelial damage caused by oral infection triggered widespread Cirl activation (Fig. 2l-r), and knocking down Cirl in ISCs and EBs or under the control of the Cirl-Gal4 promoter or Toll-8 in ECs partially reduced the proliferative response associated with this condition (Fig. 5a and S4a). While this observation might seem at odds with Cirl acting as a brake on asymmetric ISC-EB divisions, we speculated that Cirl might be required to trigger the observed switch in division mode towards symmetric ISC-ISC divisions associated with widespread damage and severe intestinal infections (Tian, Wang et al. 2017, Zhai, Boquete et al. 2017, Hu and Jasper 2019). Indeed, ISC/EB-specific Cirl depletion triggered an increase in the EB pool accompanied by a reduction in the ISC pool in this condition (Fig. S4b-k). To resolve division mode directly, we used twin-spot MARCM clonal analysis, which labels the two daughters of each division with distinct heritable markers (Fig. 5b). Symmetric ISC-ISC divisions accounted for approximately half of all divisions in infected control animals. Cirl depletion shifted this ratio to roughly 10% symmetric and 90% asymmetric (Fig. 5c-g), demonstrating that Cirl actively promotes the symmetric division mode required for regenerative amplification, extending its role beyond homeostatic fate restraint into the coordination of regenerative division outcomes.

**Figure 5:**
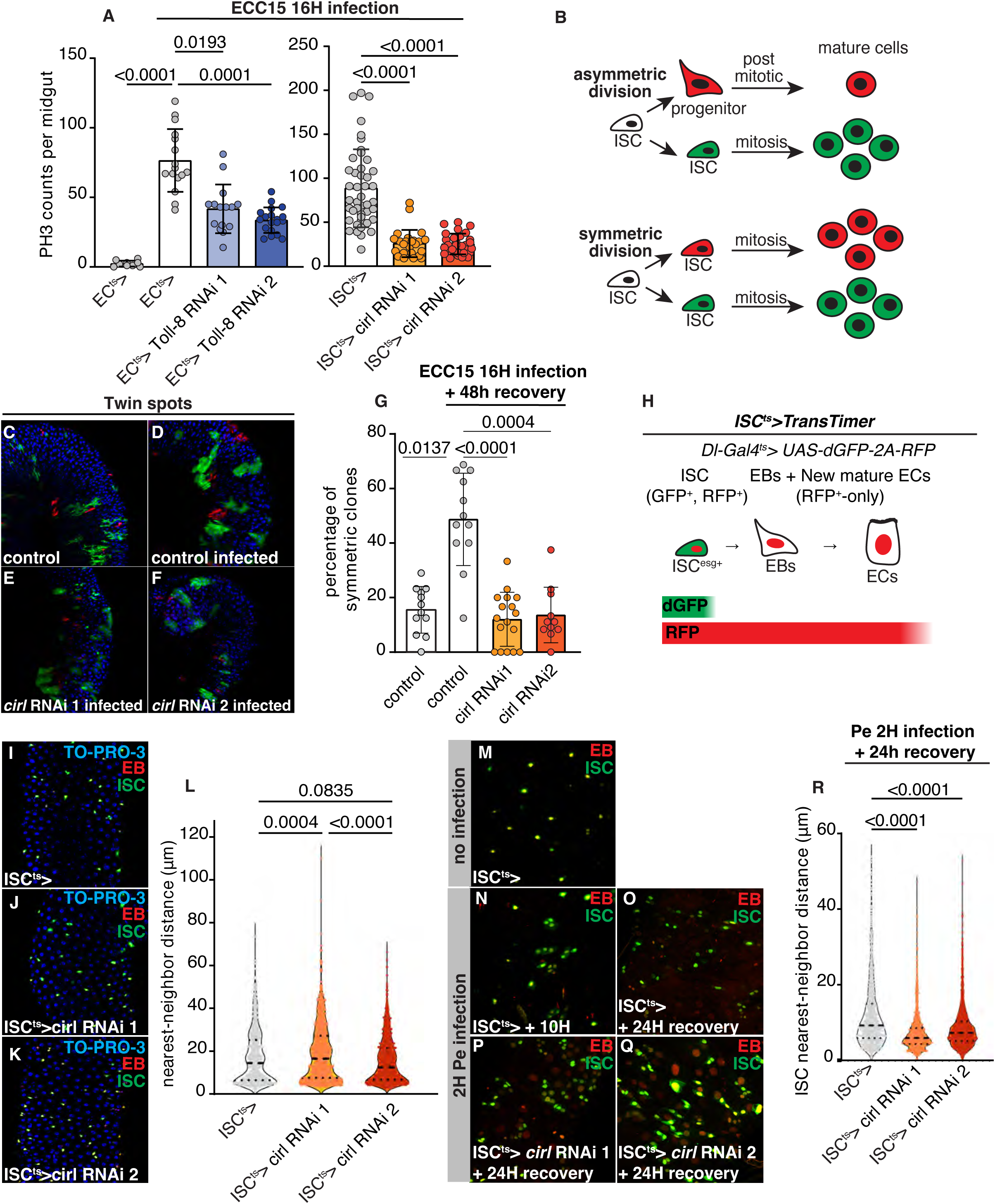
Higher-order neighbourhood sensing directs symmetric ISC divisions and tissue repatterning after injury. (a) Quantification of PH3+ cells in midguts from control flies and flies with knockdown of Cirl in ISCs (*ISC^ts^>*Cirl RNAi) or EC-specific Toll-8 depletion (*mex>*Toll-8 RNAi) following a 16-hour oral Ecc15 infection (n = 9, 16, 15, 19, 44, 26, 29). (b) Schematics of the Twin-spot MARCM (Mosaic Analysis with a Repressible Cell Marker) system. Twin-spot analyses was used to assess the effect of Cirl depletion on ISC division outcome (symmetric vs asymmetric). FLP/FRT-mediated mitotic recombination differentially labels the two sister lineages arising from a single ISC division with GFP and RFP, allowing direct determination of division outcomes and daughter-cell fates. (c-g) Cirl depletion markedly shifted division outcomes from symmetric self-renewing divisions toward asymmetric divisions during regeneration, as quantified in (g, n = 12, 12, 17, 11). (h) Schematic of the *TransTimer* system. Delta-Gal4 driven expression of short-lived GFP and a long-lived Histone(His)-2B-RFP labels ISCs in green and red, while newly generated progenitors retain only His-2B-RFP that remains stable after its expression is turned off in EBs. (i-q) Confocal images of posterior midguts with ISCs labelled in green and red (TransTimer^ts^>). (i-l) Nearest neighbour analyses show that ISC-specific Cirl depletion does not decrease ISC-ISC spacing in homeostatic conditions (l, n = 888, 1064, 1285 cells). (m-r) Confocal images of posterior midguts from control (m) and infected flies (n-q) with ISCs labelled in red and green and EBs/ECs in red alone. Acute infection induces transient ISC clustering (m-n), which resolves after 24 h of recovery (o). ISC-specific Cirl depletion impairs restoration of homeostatic ISC spacing (p-q), resulting in reduced nearest-neighbour ISC–ISC distances (r, n = 1174, 1392, 2202 cells). Statistical tests: Kruskal–Wallis with post-hoc multiple comparison analysis. Data are presented as mean values ± SD.

We tested the prediction that Cirl depletion should impair repatterning after injury without blocking the initial damage response, using localized epithelial damage. Following injury, ISCs migrate toward injured sites and assemble into dense regenerative clusters, transiently dissolving the neighbourhood relationships that sustain Cirl-Toll-8 engagement (Mackay, John et al. 2025). We first confirmed that ISC-specific Cirl depletion did not impair ISC spacing under homeostasis or migration toward damage sites (Fig. 5h-l and S4l-p), showing that higher-order signalling is dispensable for the initial injury response. In control animals, near-normal ISC spacing was restored within 24 hours of damage as clusters resolved and cells re-integrated into the epithelium (Fig. 5m-o). In Cirl-depleted animals, progenitors remained clustered at 24 hours with no evidence of spatial reorganization (Fig. 5o-r), confirming the model prediction and establishing that higher-order neighbourhood sensing provides the spatial information required to guide the transition from a regenerative to a patterned homeostatic state.

## Discussion

Many adult tissues, e.g. the respiratory epithelium, interfollicular epidermis and Drosophila midgut (Blanpain and Fuchs 2009, Tata and Rajagopal 2017, Joly and Rousset 2020, Hewitt and Lloyd 2021, O’Brien 2022), display spatial organizations that exceed simple two-state patterns, comprising multiple differentiated cell types arranged in stereotyped geometries, requiring additional organizing modules beyond classical Delta-Notch lateral inhibition. Such higher-order patterns require cells to integrate information from multiple neighbouring identities with stem cell fate decisions, allowing tissue organization to emerge from local interactions rather than a global positional signal.

Our findings support a model in which Cirl–Toll-8-mediated biomechanical signalling couples neighbourhood composition to Delta–Notch-mediated fate decisions, stabilizing tissue organization and progenitor numbers. Complementary expression of Cirl in ISCs and EBs and Toll-8 in ECs restricts receptor engagement to heterotypic interfaces, where it increases cortical myosin, restraining Delta endocytosis and Notch activation. Although each Cirl–Toll-8 interaction is pairwise, the resulting mechanical state integrates information across multiple EC contacts to modulate ISC-EB fate decisions, coupling neighbourhood composition to progenitor fate. More broadly, these findings identify adhesion GPCR-mediated mechanotransduction as a mechanism by which spatial identity is converted into cell fate.

The mechanochemical model provides a quantitative framework for this higher-order coupling. Unlike existing models based on pairwise lateral inhibition or stochastic competition (Bajpai, Chelakkot et al. 2022, Galbraith, Bocci et al. 2022), the proposed modelling framework links fate decisions to neighbourhood composition and generates stable multicellular architecture as an emergent attractor. The resulting Waddington-like landscape reveals a sharp stability boundary, predicting that modest reductions in higher-order coupling can cause a disproportionate loss of progenitor organization.

The regenerative response demonstrates that higher-order coupling is required not only to maintain tissue architecture, but also to restore it after injury. Cirl promotes the symmetric stem cell divisions that support regenerative expansion and is subsequently required for progenitors to recover their normal spatial organization. These distinct functions suggest that Cirl–Toll-8-mediated neighbourhood sensing adapts progenitor behaviour as tissue composition changes during regeneration, coordinating regenerative expansion with subsequent repatterning.

The involvement of Toll-8 extends a broader logic of identity-dependent interface recognition into adult tissue maintenance. In Drosophila, long Tolls operate as surface organizers of intercellular tension, translating differences in cell fate into mechanical asymmetries that prevent cell mixing and preserve pattern fidelity during development (Pare, Vichas et al. 2014, Pare and Zallen 2020, Umetsu 2022). A defining feature of these interfaces is the deposition and planar polarization of Myo-II at boundaries between Toll-expressing and non-expressing cells (Kolesnikov and Beckendorf 2007, Pare, Vichas et al. 2014, Lavalou, Mao et al. 2021, Tamada, Shi et al. 2021, Frey, Ernst et al. 2026). Indeed, sharp discontinuities in Toll expression within Toll-expressing domains are sensed as potential mis-specification events triggering contractile reinforcement and elimination of aberrant cells, a process referred to as interface surveillance (Frey, Ernst et al. 2026). Here, the adult epithelium appears to co-opt this interface-based recognition system to balance stability with plasticity. This suggests that the ancestral capacity of Toll receptors to discriminate “non-self” or “altered self” has been broadly repurposed beyond innate immunity as a developmental and homeostatic strategy that reinforces tissue organization through mechanically active interfaces.

Whether equivalent higher-order interactions operate in vertebrate tissues remains an open question, but structural and functional parallels provide a possible basis. Mammalian Toll-like receptors (TLRs) contain extracellular leucine-rich repeat domains with structural similarity to FLRT proteins (Dolan, Walshe et al. 2007), established heterophilic ligands of Latrophilins (O’Sullivan, de Wit et al. 2012, Boucard, Maxeiner et al. 2014, Sando, Jiang et al. 2019, Del Toro, Carrasquero-Ordaz et al. 2020, Li, Xie et al. 2020) and can modulate actomyosin dynamics independently of canonical NF-κB signalling (Peterson, Balogh Sivars et al. 2023). Whether a Latrophilin–TLR axis implements analogous neighbourhood sensing in vertebrate regenerative epithelia remains to be established, but the structural parallels and shared mechanical outputs provide a testable framework.

Our findings identify receptor-mediated mechanotransduction as a mechanism that couples neighbourhood composition to stem cell fate, enabling higher-order spatial organization to emerge from local interactions. Such mechanisms may represent a general solution to the challenge of maintaining complex cellular patterns in dynamic, regenerative tissues. Understanding how this coupling is calibrated, how it fails in disease and whether it can be re-engaged to restore complex tissue architecture will be important directions for future work.

## Acknowledgements

J.C. and D.S.A. are funded by H2020 European Research Council grant number 803630, Novo Nordisk Foundation grant number NNF180C0033920, and Danish Research Council grant number 4283-00049B. A.J. was funded by the Horizon-MSCA grant number 101109581. We thank the Carlsberg foundation for equipment grants CF19-0353 and CF23-1302. A. D. acknowledges funding from the Novo Nordisk Foundation (grant No. NNF18SA0035142 and NERD grant No. NNF21OC0068687), Villum Fonden (Grant no. 29476), and the European Union (ERC, PhysCoMeT, 101041418). Views and opinions expressed are, however, those of the authors only and do not necessarily reflect those of the European Union or the European Research Council. Neither the European Union nor the granting authority can be held responsible for them.

## Author contributions

A.J., T.M., A.D., J.C., and D.S.A. designed the research, A.J. and T.M. conducted most experiments for the manuscript with the support of D.M. and E.C., D.M., A.J., J.C., and D.S.A. performed the genetic screen that formed the basis for the Cirl-Toll-8 project, A.J., T.M., J.C., A.D., and D.S.A. analyzed the data, J.C., A.D., and D.S.A. supervised the project, and D.S.A. and A.D. wrote the manuscript.

## Materials and methods Fly stocks and husbandry

### Husbandry

For practical reasons, mated female Drosophila melanogaster was used exclusively in all experiments. Adult flies and crosses were maintained in environmentally controlled incubators at 18°C, 25°C or 29°C with 60% relative humidity under a standard 12-hour light/dark cycle. Animals were reared on a standard cornmeal-yeast-sucrose diet consisting of 82 g/L cornmeal, 60 g/L sucrose, 34 g/L baker’s yeast, and 8 g/L agar. The medium was supplemented with 4.8 mL/L propionic acid and 1.6 g/L methyl-4-hydroxybenzoate to inhibit microbial growth. To maintain optimal health and colony hygiene, flies were transferred to fresh medium every 48 hours.

To achieve spatiotemporal control of transgene expression while avoiding developmental abnormalities, we utilized the temperature-sensitive TARGET system (Mcguire et al., Science, 2003) (tub:Gal80^ts^). Flies were raised at a restrictive temperature of 18°C throughout embryonic, larval, and pupal development. Following eclosion, adults were maintained at 18°C for an additional four days to ensure complete maturation of the adult digestive system. To initiate Gal4-mediated transgene expression in a cell-type-specific manner, the adult flies were subsequently shifted to a permissive temperature of 29°C. Unless otherwise specified in the text, this period of UAS induction was maintained for 5 to 7 days before experimental analysis.

### Drosophila Stocks and Genetics

We are grateful to the *Drosophila* research community for generously providing several of the transgenic lines used in this study. The following stocks were graciously gifted: Esg-ReDDM – UAS CD8::GFP, esg-Gal4/CyO; UAS H_2_B::RFP, tub-Gal80^ts^/TM6b (Maria Dominguez, IN, Spain), endo ECad::3xGFP (Yohanns Bellache, Institut Curie, France), UAS:transtimer – UAS dGFP::2A::RFP and Su(H)Gbe:eRFP (Norbert Perrimon, Harvard Medical School, USA), ISC^ts^ – esg-Gal4, UAS YFP; Su(H)Gbe:Gal80, tub-Gal80ts (Heinrich Jasper, Genentech, USA), EC Rapport – esg-LexA, lexAop-mCD8GFP, lexAop-H_2_B::mCherry, tub-Gal80^ts^/CyO; mex:Gal4 (Tobias Reiff, Heinrich Heine University Düsseldorf, Germany), Toll-8::YFP (Jennifer Zallen, Sloan Kettering Institute, USA), UAS Toll-8::HA (Konrad Basler, University of Zurich, Switzerland), CFS - w;esg-Gal4,UAS CFP,Su(H)Gbe-nls::GFP; tub-Gal80^ts^ (Lucy O’brien, Stanford University, USA), Sqh::GFP [KI] (Magali Suzanne, CBI Toulouse), Dl::GFP (François Schweisguth, Institut Pasteur, France), esg-Gal4, UAS myr::Tomato, tub-Gal80^ts^/CyO; Pros::GFP/TM6b (Zheng Guo, HUST, China), and Toll-2 Gal4, Toll-6 Gal4, and Toll-7 Gal4 (Alicia Hidalgo, University of Birmingham, UK).

UAS cirl gRNA (97786), UAS cirl RNAi 1 (34821), UAS cirl RNAi 2 (27524), cirl-Gal4 (CRIMIC) (91479), UAS lifeAct::Ruby (35545), Sqh::mScarlet [KI] (94929), Delta-Gal4 (Trojan) (77753), hs-FLP (7), UAS mCD8GFP, UAS CD2 RNAi, FRT40A/CyO; TM3/TM6 (56185), UAS- CD2::RFP, UAS GFP RNAi, FRT40A/CyO; TM3/TM6 (56184), UAS Toll-8 RNAi 1 (28519), act-FRT>>CD2>>FRT-Gal4 (4779), UAS SqhDD/CyO; NRE:EGFP (30727), NRE:EGFP (30728), TM2/TM6 (600573), UAS luciferase RNAi (TRiP) (31603), UAS mCD8::RFP, LexAop mCD8::GFP; CoinFLP-LexA::GAD.Gal4 (58754), UAS Toll- 8^FL^::EGFP/CyO (92990), and UAS:Toll-8^ΔCyto^::EGFP/CyO (92991) were acquired from the Bloomington Drosophila Stock Center (BDSC). UAS sqh RNAi 1 (7917), UAS sqhRNAi 2 (7916), UAS Toll-8 RNAi 2 (13549) were acquired from the Vienna Drosophila Resource Center (VDRC).

### Fly strains generated in this study

To generate the Cirl mechanosensory lines (UAS Cirl::NRS::LexA, UAS Cirl ΔGPS::NRS::LexA, and UAS Cirl NRSΔS3::LexA), Cirl NRS::LexA, Cirl ΔGPS::NRS::LexA, and Cirl NRSΔS3::LexA coding sequences were PCR-amplified from pAD4, pAD3, and pAD29 plasmids (kindly gifted by Tobias Langenhan, Leipzig University, Germany (Scholz et al., Nature, 2023)), respectively, and cloned into the pENTR/D-TOPO vector using the following sense primer CAC CAT GGA GAC AGA CAC ACT CC and antisense primer TTA TTT TAG AAC TCG AAC CTC GAT GAA CAT G. Cirl NRS::LexA, Cirl ΔGPS NRS::LexA, and Cirl NRSΔS3::LexA coding sequences were subcloned into pUASg.attB and pUASattB Cirl NRS::LexA, pUASattB CirlΔGPS::NRS::LexA, and pUASattB Cirl NRSΔS3::LexA constructs were introduced into the germ line by injections in the presence of the PhiC31 integrase and inserted in the 86F8-landing site on the 3R chromosome (BDSC-BL24749, BestGene).

### Bacterial Culture and Oral Infection Assays

Oral infections were performed using either *Erwinia carotovora carotovora* 15 (Ecc15) or *Pseudomonas entomophila* (Pe). Bacterial cultures were prepared overnight in conical flasks containing Luria-Bertani (LB) broth. Ecc15 cultures were inoculated from single colonies and incubated at 29°C. Pe cultures were inoculated directly from −80°C glycerol stocks and grown at 30°C in LB broth supplemented with rifampicin. Following overnight growth, the optical density OD600 of each culture was measured. The cultures were centrifuged, and the resulting bacterial pellets were resuspended in a 5% sucrose solution to achieve specific infectious doses. The final bacterial suspensions were adjusted to an OD600 of 100 for assays measuring phosphohistone H3 (PH3+) in response to Pe infection, and an OD600 of 200 for all other experiments.

To ensure efficient and consistent consumption of the bacteria, adult flies were subjected to a brief starvation period prior to infection. Flies were sorted into cohorts of 10 per empty vial and starved for 2 to 3 hours. During the starvation period, infection vials were prepared by placing Whatman filter paper disks on the surface of standard fly food medium. Each disk was inoculated with 50 µl of the concentrated bacteria solution. The starved flies were subsequently transferred into these prepared vials and allowed to feed on the bacterial medium for the indicated experimental time points.

### Dissections and Immunohistochemistry

Adult midguts were dissected in phosphate-buffered saline (PBS) and immediately transferred to a fixative solution of 4% paraformaldehyde in PBS for 1 hour at room temperature. Following fixation, tissues were washed twice in standard PBS, followed by two 15-minute washes in PBS containing 0.1% Triton X-100 (PBS-T) under gentle agitation. Samples harboring endogenously fluorescent reporters that did not require further antibody staining were subjected to a final PBS wash and immediately mounted on microscopy slides. For samples requiring immunohistochemistry, fixed midguts were incubated in a blocking solution consisting of 10% fetal bovine serum in PBS-T for 2 hours at room temperature, then incubated with primary antibodies diluted in blocking solution overnight at 4°C. The following day, midguts were washed three times for 15 minutes each in PBS-T, then incubated with secondary antibodies for either 3 hours at room temperature or overnight at 4°C. After secondary antibody incubation, tissues were washed twice for 15 minutes in PBS-T, followed by a final single wash in PBS. The primary antibody used is rabbit anti-phosphohistone H3 (1:1000; Millipore 06-570), detected with Alexa Fluor 488-conjugated goat anti-rabbit (1:1000; Thermo Fisher A-11008) secondary antibody. All prepared midguts were mounted on glass slides using Vectashield mounting medium, with or without DAPI for nuclear counterstaining. In cell fate sensor experiments utilizing CFP as a fluorophore, TO-PRO-3 iodide (Catalog # T3605, Thermo Fisher Scientific) was used as a generic nuclear label. To preserve three-dimensional tissue morphology, samples were mounted using 0.12 mm SecureSeal spacers (Grace Bio-Labs), which were omitted exclusively for experiments assessing the total number of mitotic cells per gut to facilitate whole-gut cell counting.

### Confocal Microscopy and Image Processing

#### Fixed samples

Images of fixed gut tissues were acquired using two systems: an inverted Zeiss LSM-900 confocal microscope controlled by Zen Blue software (utilizing 5x and 20x objectives), and an Andor Dragonfly 200 spinning disk confocal microscope equipped with a Leica 40x oil immersion objective. For volumetric analysis, Z-stacks were acquired at 1 μm intervals to capture the entire depth of the intestinal epithelium, spanning from the basal surface to the lumen. For both qualitative observation and subsequent quantification, imaging was localized to the posterior gut. Acquisitions consisted of either the complete R4 region or representative single-tile scans within the R4BC subregion. Post-acquisition image processing and formatting were performed using Fiji (ImageJ) and Adobe Photoshop.

#### Live gut explants

To preserve the membrane localization of fluorescently labeled proteins sensitive to chemical fixation (e.g., Delta, E-Cad, and Myo-II), gut explants were processed using a live-cell imaging protocol. Whole midguts were dissected in modified Schneider’s medium, as previously described (Marchetti et al., 2022). Following dissection, the midguts were mounted in 35 mm coverslip-bottom imaging dishes (Ibidi, Cat# 80136) and embedded in 0.5% low-melting-point agarose prepared in modified Schneider’s medium. Prior to imaging, 100 μL of modified Schneider’s medium supplemented with the calcium channel blocker isradipine (10 μg/mL; Sigma-Aldrich, Cat# I6658) was added to the dish.

Live imaging was performed using a Leica DMi8 inverted microscope equipped with an Andor Dragonfly 200 spinning disk confocal scanner. Acquisitions were performed using a 40x oil immersion objective (NA 1.3) and excited with 488 nm and 561 nm laser lines. Confocal Z-stacks were acquired at 1 μm intervals. Post-acquisition image processing and formatting were performed using Fiji (ImageJ) and Adobe Photoshop.

### TWINSPOT-MARCM clones

To trace the mode of stem cell division and simultaneously knock down Cirl, twin-spot MARCM crosses were established using a driver stock carrying act-FRT>>CD2>>FRT-Gal4, hs-FLP, fluorophores, and the respective RNAi construct. Crosses and progeny were maintained at 18°C to prevent spontaneous FLP-mediated recombination. At 18°C, the FRT-flanked CD2 cassette effectively blocks Gal4-mediated expression, keeping the RNAi construct inactive, helping to prevent RNAi expression during development.

Newly eclosed F1 adult flies were maintained at 18°C for 5 days to allow the digestive epithelium to fully mature. To induce clones, flies were subjected to a single 30-minute heat shock at 37°C. Heat shock induction of the FLP recombinase excises the CD2 cassette, activating Gal4-mediated expression. When this recombination occurs in a mitotic intestinal stem cell (ISC), it generates one RFP-marked and one GFP-marked daughter cell, with both clonal lineages expressing the RNAi construct. Following heat shock, induced flies were returned to 18°C for 10 days to allow sufficient time for RNAi-mediated reduction of Cirl protein levels. Flies were then subjected to bacterial infection using *Ecc15* for 16 hours, transferred to fresh food, and allowed to recover for 2 days. Whole posterior midguts were subsequently dissected, fixed, and imaged entirely using the Andor Dragonfly 200 spinning disk confocal microscope.

### ISC Cluster Analysis

To evaluate ISC clustering, adult flies were subjected to an oral infection paradigm. Prior to infection, flies were starved in empty vials for 2–3 hours. Following starvation, flies were fed a 5% sucrose solution supplemented with *Pe* at a final OD600 of 200. This infection phase was maintained for 2 hours. The flies were then transferred back to standard medium for a 4-hour recovery period prior to dissection, fixation, and subsequent imaging.

### Image Analysis and Quantification

#### General Image Processing and Cell Counting

All image analyses and quantifications were performed using the open-source software Fiji (ImageJ). For quantitative measurements of specific cell types, Z-stacks acquired with a 20x objective in the posterior midgut R4BC region were converted to maximum intensity projections (MIPs). Cell numbers were manually annotated using the Fiji CellCounter plugin. Total cell counts were normalized to the measured epithelial area or total cells in the area, with cell density consistently defined and reported as cells per 10,000 µm² or ratio of cell type respectively.

#### Spatial Distribution and Topological Patterning Analysis

To evaluate the spatial distribution and topological patterning of ISCs, an intercellular ’hop distance’ metric was employed. For this analysis, a linear trajectory was drawn connecting the centroids of adjacent ISCs. The topological distance between cells was quantified by counting the number of EC plasma membrane boundaries intersected by this connecting line. The total number of intervening ECs separating the two ISCs was subsequently calculated by subtracting one from the total number of intersected boundaries.

To infer the biological rules governing ISC spatial localization, the calculated hop distances were aggregated and plotted as frequency histograms. Tissue patterning was classified based on the resulting histogram profiles: a dominant frequency peak at a distance of 0 to 1 intervening cells was classified as spatial clustering; a peak at 2 to 3 intervening cells was indicative of a specific separation radius between ISCs.

#### Cell Segmentation and ISC Clustering

To quantify ISC clustering, Z-stacks of imaged midguts were converted to MIPs and segmented using Cellpose (utilizing the cpSAM model). The automated Cellpose-generated masks were manually inspected and corrected to ensure absolute segmentation accuracy. The finalized masks were imported into Fiji, where spatial clustering was analyzed using the SSIDC (Spatial Statistics based on Inter-Distance Clustering) cluster indicator within the BioVoxxel toolbox. A maximum intercellular distance (epsilon) of 5 pixels (corresponding to 2 µm) was applied to define clustered cells.

#### Stem Cell Morphometric Analysis

To analyze ISC morphometrics following cirl knockdown, single-cell binary masks were generated using Cellpose. These masks were processed using the Extended Particle Analyzer within the Fiji BioVoxxel toolbox to extract key shape descriptors like area.

#### Fluorescence Intensity Measurement

To quantify the mean fluorescence intensity of membrane-localized Dl::GFP and MyoII::GFP, Z-stacks were restricted to a 5 µm volume starting from the basal-most surface of the intestinal epithelium and converted to MIPs. The boundaries of ISCs and EBs were segmented using Cellpose, and the resulting outlines were saved as ROIs. These ROIs were subsequently used to measure the mean fluorescence intensity of the membrane-localized proteins.

For the quantification of MyoII::GFP following Toll-8 knockdown in enterocytes, a modified approach was required due to the absence of a distinct ISC/EB fluorescent marker. In these instances, the mean fluorescence intensity of Myo-II at the cell membrane was manually quantified using the Segmented Line tool in Fiji.

#### Scoring of Symmetric and Asymmetric Divisions

To ensure accuracy, a low density of twin-spot induction was utilized, allowing for the reliable identification of daughter pair fates even if the marked clones were not perfectly juxtaposed. Only initial labeling divisions were scored, and analysis was restricted to twin spots containing a total of ≥3 cells (i.e., ≥2 cells of one color and ≥1 cell of the other color).

Fate outcomes were categorized based on cellular composition:

##### Asymmetric outcomes

Twin spots containing exactly 1 cell of one color and ≥2 cells of the other color. In these instances, the uniquely labeled cell had nearly always differentiated into a polyploid enterocyte by the time of analysis, while the stem cell continued to generate daughters.

##### Symmetric outcomes

Twin spots where both the RFP and GFP clones contained ≥2 cells, indicating both initial daughter cells retained proliferative capacity. In these cases, each colored twin spot nearly always contained at least one diploid cell

#### Nearest Neighbor Distance Analysis

To quantify the spatial proximity of intestinal stem cells (ISCs), a nearest-neighbor distance (NND) analysis was performed. Following the automated segmentation of ISCs and generation of binary masks using Cellpose, the X and Y coordinates of each cell centroid were extracted within Fiji. The complete matrix of Euclidean distances between all individual ISCs within each midgut was subsequently calculated. For each distinct ISC, the minimum calculated Euclidean distance to an adjacent ISC was identified using a custom Excel macro, with this minimum value *defined as the NND*.

### Mechanochemical Delta–Notch vertex model of stem-cell–pair patterning

#### 1. Tissue mechanics

Cells tile the surface as polygons whose vertices *r* evolve by overdamped relaxation of the vertex energy

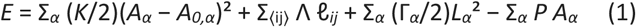

with cell area *A_α_*, perimeter *L_α_*, edge length ℓ*_ij_*, area stiffness *K*, line tension Λ (held constant, with weak Ornstein–Uhlenbeck noise), cortical contractility Γ*_α_*, and pressure *P* (zero on the periodic torus). Vertices follow

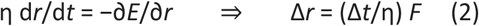

integrated explicitly with viscosity η and step Δ*t*; neighbour exchange (T1) occurs below edge length *t_1_*.

Two geometries are used. (i) A periodic torus with area-conserving box expansion: the box is rescaled each step so that box area = *N*·*a_ref_* (count mode), holding the mean cell area constant as the cell number *N* grows. (ii) A free-boundary disk (round tissue) with no box expansion, whose rim is stabilised by a boundary line tension Λ_bd_; the colony grows outward as cells divide.

#### 2. EC-junction contractility and the mechanosensor gain g

A single per-cell field couples mechanics to signalling. Cortical myosin is recruited on junctions that separate a signalling cell from an EC neighbour,

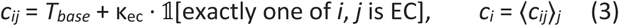

The mechanosensor (Cirl) transduces this field with gain *g*, giving an effective contractility 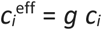 which sets both the mechanical contractility and the signalling throttle:

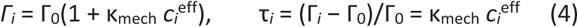

where τ*_i_* is a dimensionless effective cortical-tension index. Setting *g* = 1 recovers the wild type (Control); *g* = 0 is the full Cirl knockout, which simultaneously removes the signalling throttle and relaxes Γ*_i_* to baseline.

#### 3. Delta–Notch signalling with tension-throttled uptake

Each non-EC cell carries Delta *D_i_* and Notch *N_i_* (ECs are frozen at *D* = *N* = 0). Trans-activation by neighbour Delta is throttled by the local cortical tension:

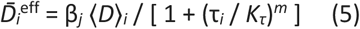

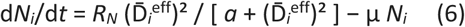

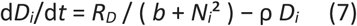

#### 4. Cell fate and proliferation

Fate is a monotone, irreversible lineage: ISC → EB → EC. An EB commits to EC when its Notch exceeds a threshold, *N_i_* > *N*_EC_. An ISC divides when its division signal exceeds *D*_div_,

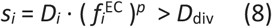

where 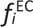 is the EC fraction among cell *i*’s neighbours and *p* is the EC-contact power. The EC-contact power (*p* = 2) is itself mechanosensor-dependent and is removed in the Cirl-knockout (*p* = 0).

Both fate transitions are **stochastic**: once a cell satisfies its signal gate it commits with probability *r* Δ*t* per step — hazard rate *r_div_* for ISC division and *r_ec_* for EB→EC commitment.

#### 5. Initial condition and order parameter

Simulations start from a confluent, uniform Delta-high tissue (all ISC, low Notch, small per-cell noise). The pattern is quantified by the isolated-pair order parameter

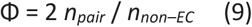

where an *isolated pair* is one ISC whose only signalling (non-EC) neighbour is a single EB, that EB’s only signalling neighbour being the same ISC, with all remaining neighbours EC. The steady-state value is the time average of Φ over the plateau (*t* ≥ 12×10³ units).

#### 6. Parameters

**Table S1.** | Mechanics. Vertex-model parameters.

| Symbol | Meaning | Value |
| --- | --- | --- |
| $K$ | area stiffness | 1.0 |
| $\Lambda$ | line tension (constant) | -0.924 |
| $\Gamma_0$ | baseline cortical contractility | 0.14 |
| $\eta$ | vertex viscosity | 0.02 |
| $\Delta t$ | integration time step | 0.001 |
| $t_{1\text{eps}}$ | T1 transition length | 0.01 |
| $\tau_{\text{ou}} / \sigma_{\text{ou}}$ | $\Lambda$ Ornstein–Uhlenbeck time / amplitude | 0.128 / 0.02 |
| $\Lambda_{\text{bd}}$ | boundary line tension (disk) | 0.5 |
| $\tau_{\text{life}}$ | cell-cycle lifespan | 3.5 |

**Table S2.**
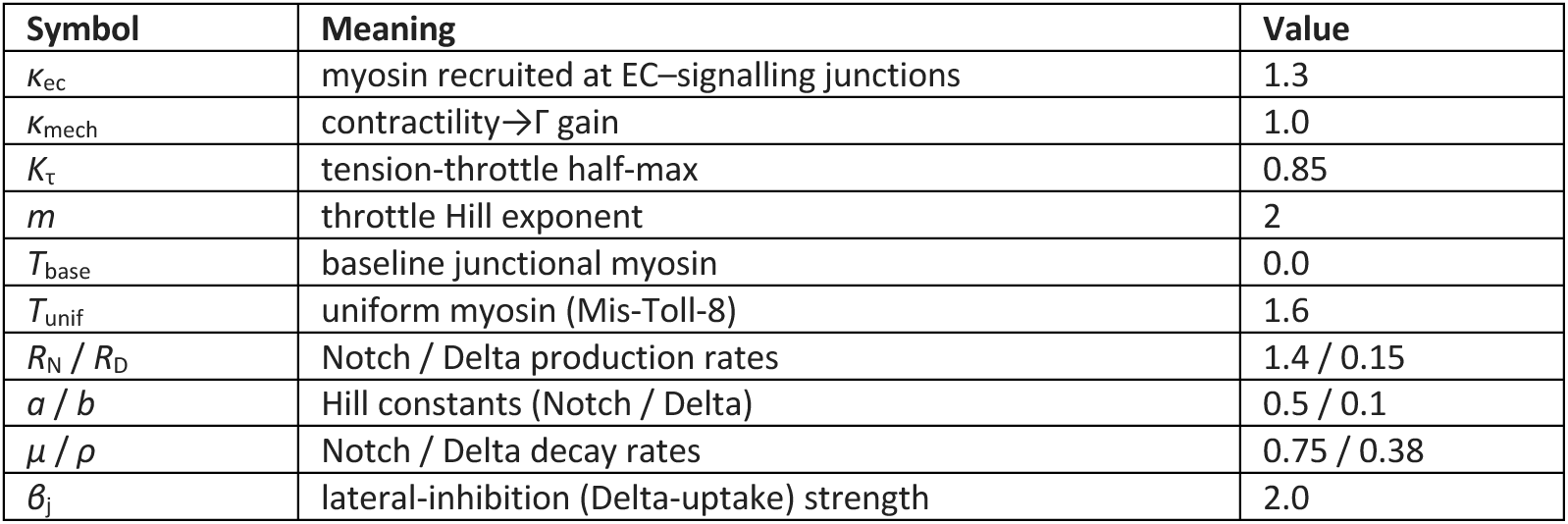
| Mechanosensor and Delta–Notch reaction.

**Table S3.** | Fate and proliferation. Fate transitions are stochastic.

| Symbol | Meaning | Value |
| --- | --- | --- |
| $N_{\text{EC}}$ | Notch threshold for EB→EC commitment | 0.85 |
| $D_{\text{div}}$ | Delta-signal threshold for ISC division | 2.5 |
| $p$ | EC-contact power of the division gate | 2 (Control) / 0 (KO) |
| $r_{\text{div}}$ | stochastic ISC-division hazard rate | 0.5 |
| $r_{\text{ec}}$ | stochastic EB→EC hazard rate | 1.0 |

#### 7. Numerical implementation and experiment settings

Equations (1–2) are integrated explicitly (Δ*t* = 10⁻³) with the Delta–Notch ODEs (5–7) advanced on the same step; times are reported in 10³ simulation units. Runs extend to *t* = 16 with snapshots every 200 steps. The baseline runs use a 10×10 hexagonal torus (100 cells, count-mode box expansion, seed 41); the self-organisation runs use a free-boundary Voronoi disk of 40 cells, reporting Φ as mean ± s.d. over six independent seeds. Control and Cirl-KO share every parameter and differ only in *g* and *p*. The model is implemented in Python (NumPy/SciPy).

## Statistical Analysis

All statistical analyses and data visualizations were performed using GraphPad Prism software. Prior to analysis, all datasets were independently assessed for normal distribution using the Shapiro-Wilk normality test. The selection of statistical tests was determined by the normality of the data distribution and the number of experimental groups being compared:

## Two-group comparisons

Normally distributed datasets were analyzed using an unpaired Student’s t-test. Datasets that did not pass the normality test were analyzed using the Mann-Whitney U test.

## Multiple-group comparisons (≥3 groups)

Normally distributed datasets were analyzed using a one-way analysis of variance (ANOVA) followed by Tukey’s multiple comparisons test. Non-normally distributed datasets were analyzed using the Kruskal-Wallis test followed by Dunn’s multiple comparisons test.

In graphical representations, error bars generally denote the standard deviation (SD) when the sample size is <50 and standard error mean (SEM) when the sample size is large.

## Movie 1. Higher-order mechanosensing stabilizes progenitor organization

Vertex-model simulations of tissue organization with intact mechanosensing (control, g=1) or complete loss of mechanosensing (Cirl KO, g=0). Control tissues maintain regularly distributed ISC–EB pairs, whereas loss of coupling leads to progressive disruption of progenitor organization and continued cell production. Delta^hi^ ISCs are shown in red, Notch^hi^ EBs in green, initial ECs in white, and newly generated ECs in grey.

## Movie 2. Higher-order mechanosensing promotes tissue repatterning

Vertex-model simulation of tissue regeneration with intact Cirl–Toll-8 mechanosensing. Starting from a progenitor-enriched state, the tissue progressively restores a spatially dispersed pattern of Delta^hi^ ISCs and Notch^hi^ EBs surrounded by ECs.

**Figure S1.**
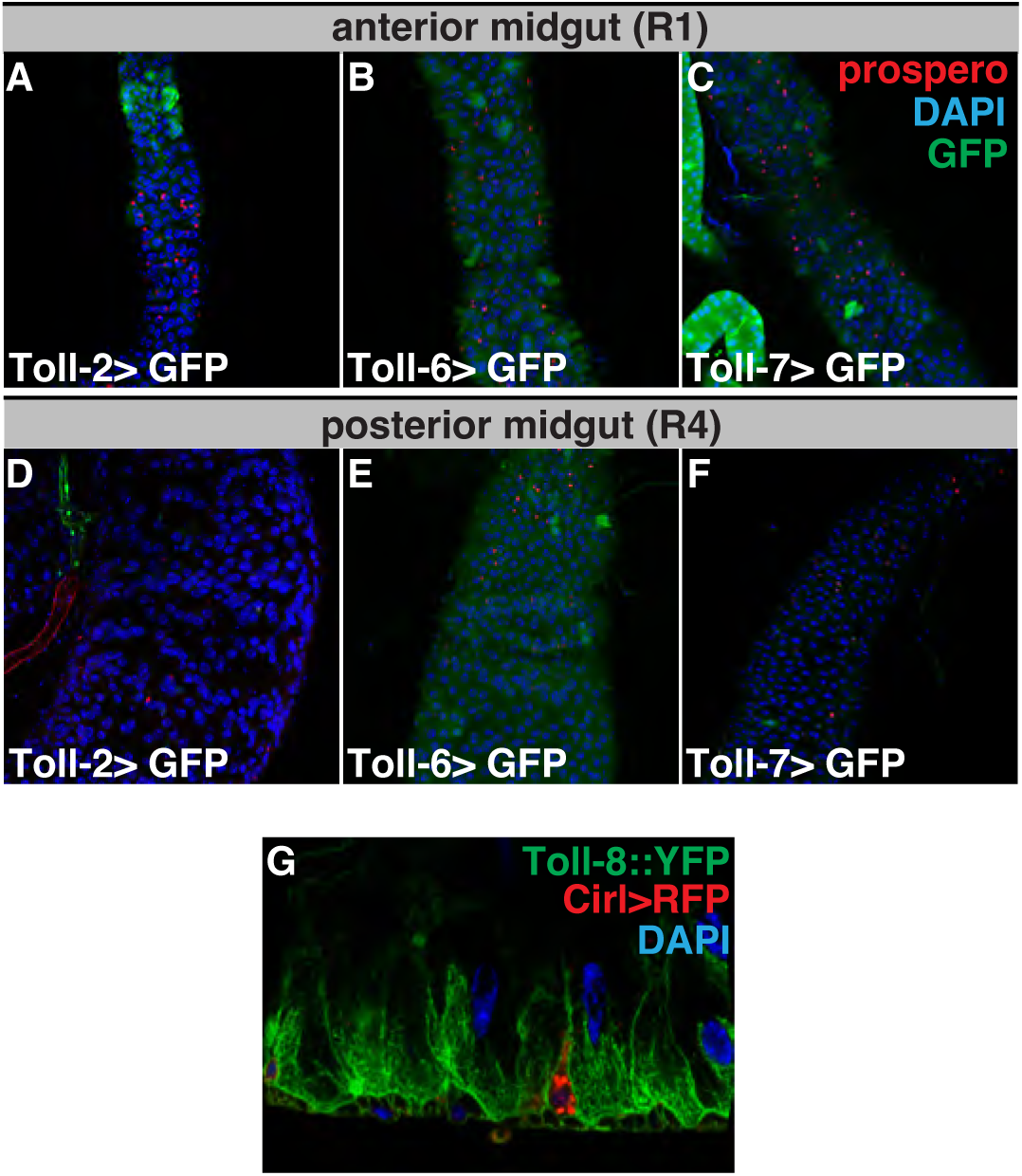
Toll-2, Toll-6 and Toll-7 are not expressed in the posterior midgut. (a-f) Confocal images of anterior (a-c) and posterior (d-f) midguts showing the expression patterns of Toll-2 (Toll-2>GFP), Toll-6 (Toll-2>GFP), and Toll-7 (Toll-2>GFP; in green). Although all three receptors are sporadically expressed in ECs of the anterior midgut, none are detected in the posterior midgut. (g) Confocal images of posterior midguts from flies expressing *cirl>UAS-RFP* together with endogenous Toll-8-YFP reveal complementary expression patterns, with Cirl restricted to ISCs/EBs and Toll-8 to ECs.

**Figure S2.**
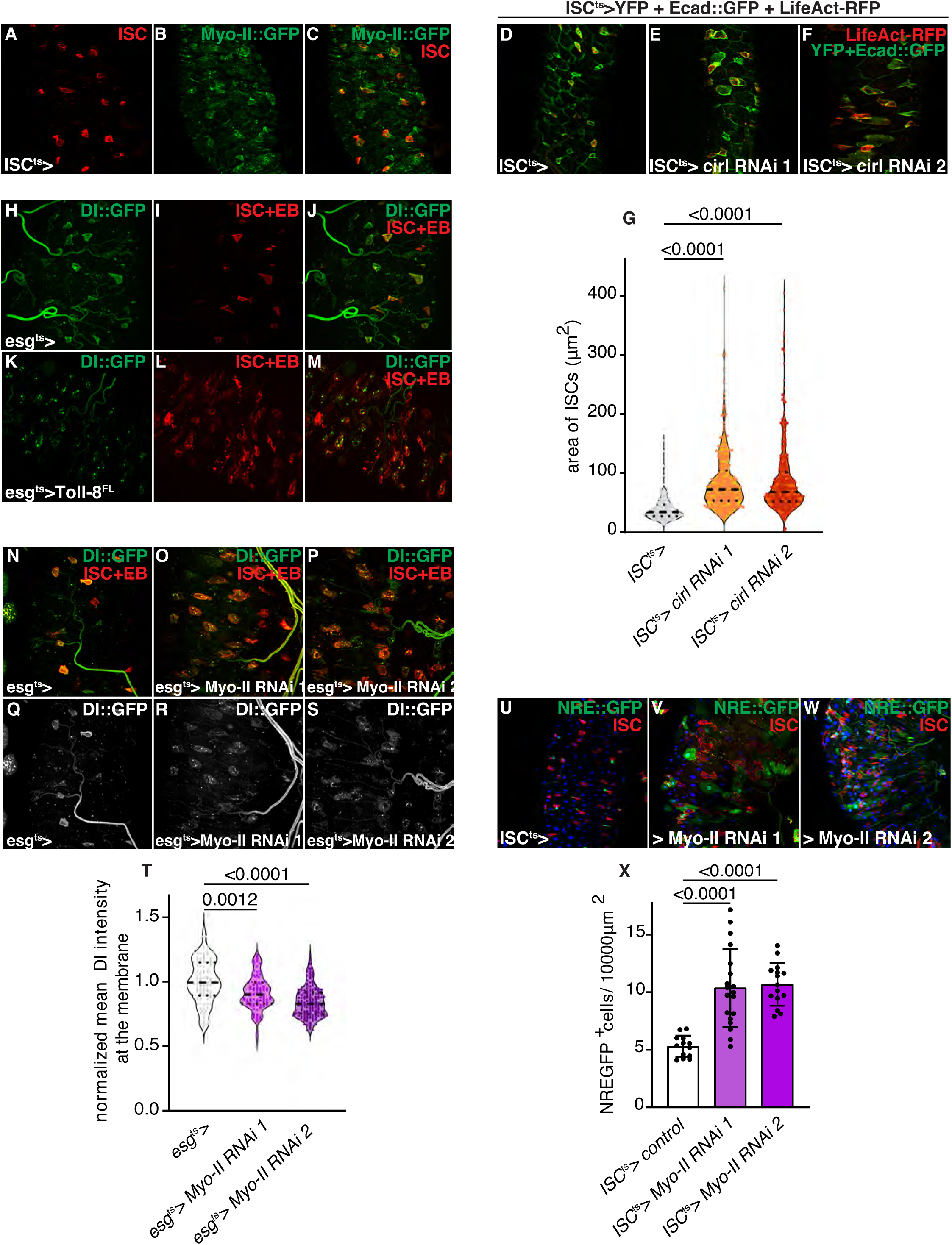
Notch activity inversely correlates with cellular tension. (a-c) Confocal images of posterior midguts dissected from flies expressing endogenous Myo-II::GFP and ISCs labelled in red (ISC^ts^>RFP) show elevated cortical Myo-II in ISCs and their EB neighbours. (d-g) Confocal images of adult posterior midguts with ISC-specific Cirl depletion reveal an increase in ISC area, consistent with reduced cortical tension, as quantified in (g, n = 199, 267, 307 cells). (h-s) Confocal images of posterior midguts expressing endogenous Delta::GFP. (h-m) ISC-EB-specific overexpression of Toll-8^FL^ increases Delta-GFP internalization into intracellular vesicles. (n-t) Notch activity inversely correlates with cellular tension. (n-s) ISC-EB-specific depletion of Myo-II enhances Delta::GFP internalization, resulting in decreased membrane-associated Dl, as quantified in (t, (n = 99, 92, 157 cells). (u-w) Confocal images of posterior midguts dissected from flies expressing the Notch activity reporter (NRE::GFP). ISC-EB-specific depletion of Myo-II increased Notch reporter activity as quantified in (x, n = 13, 15, 15). Statistical tests: Kruskal–Wallis with post-hoc multiple comparison analysis. Data are presented as mean values ± SD.

**Figure S3.**
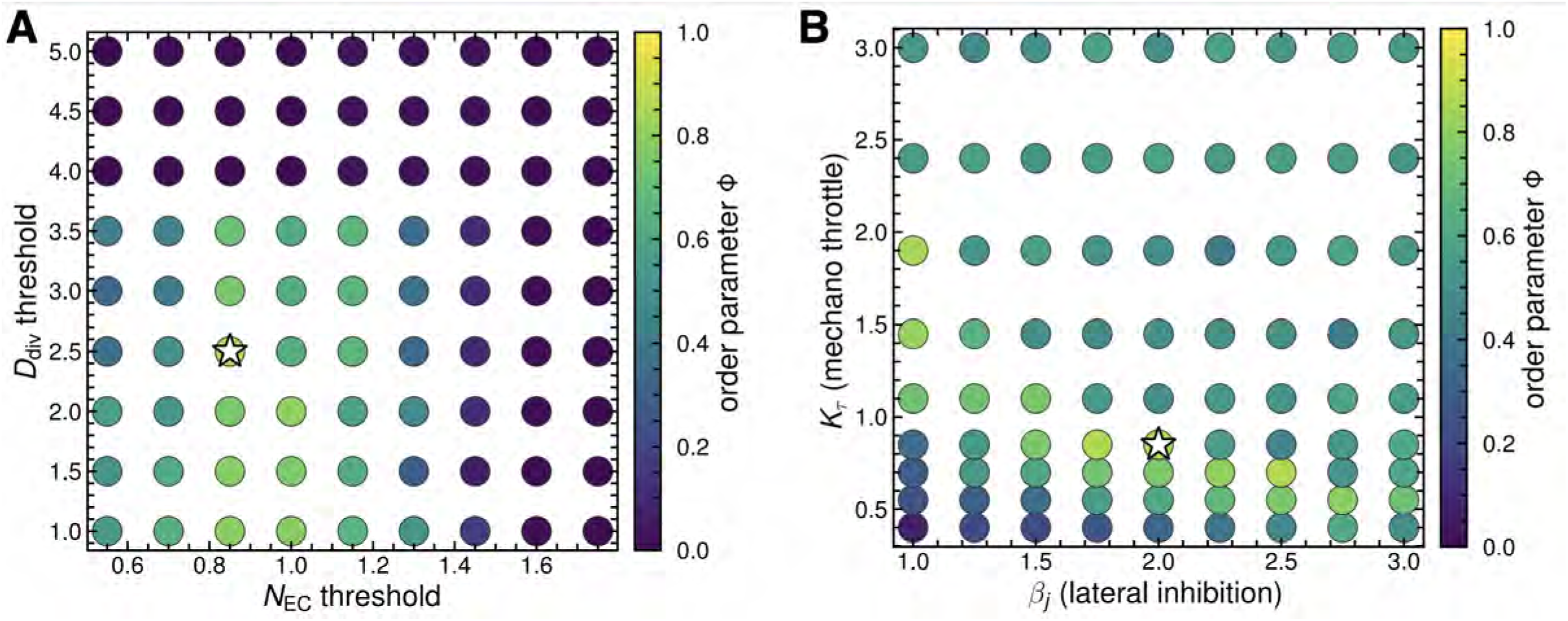
Sensitivity of tissue organization to model parameters. (a) Sensitivity analysis of the steady-state order parameter, φ, across combinations of the EC differentiation threshold, N_EC_ threshold, and the ISC division threshold, D_div_ threshold. (b) Sensitivity analysis of φ across combinations of the lateral-inhibition strength, β_i_, and the mechanosensing threshold, K_T_. In both panels, circle colour indicates the resulting order parameter, with higher values corresponding to more ordered tissue organization. Stars mark the parameter combinations used in the simulations presented in the main text.

**Figure S4.**
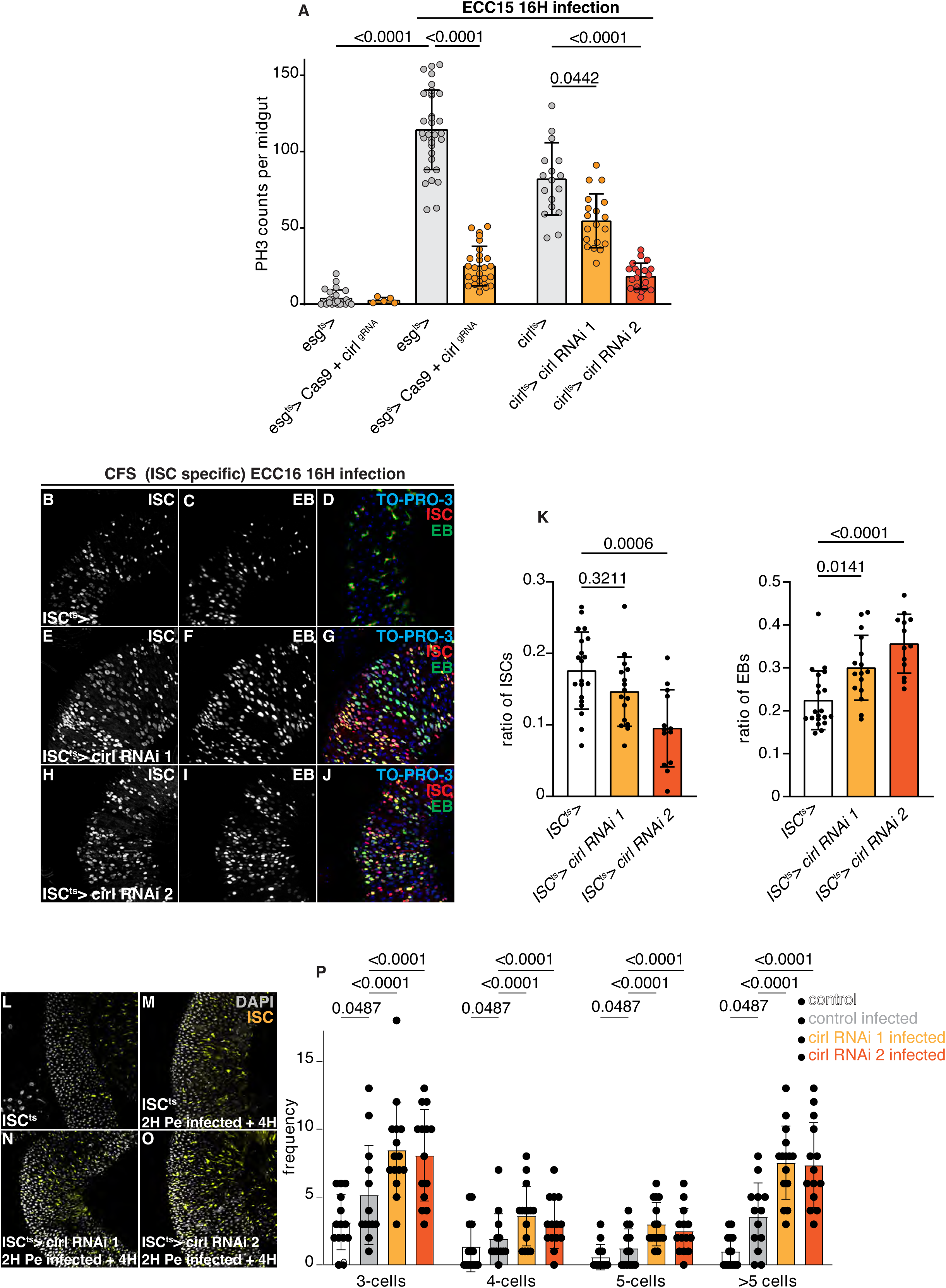
ISC-specific Cirl depletion reduces ISC numbers and expands the EB pool following oral infection. (a) Quantification of PH3+ cells in midguts from control flies and flies with knockdown of Cirl in ISCs and EBs (*esg^ts^*>Cirl RNAi) or Cirl-expressing cells (*cirl>*Cirl RNAi) following a 16-hour oral Ecc15 infection (n = 24, 5, 33, 27, 18, 19, 20). (b-j) Representative confocal images of posterior midguts showing that ISC-specific (CFS^ISCts^>) Cirl depletion increases the EB population (in green) and reduces the ISC pool (in red) as quantified in k (n = 20, 16, 13). (l-o) Confocal images of posterior midguts with ISCs labelled in green dissected from uninfected flies (l, control) or flies exposed to P.e. for 2 hours (m-o). Brief exposure to P.e. infection for 2 hours triggers the formation of ISC clusters 6 hours post infection (m), and this is not prevented by ISC-specific Cirl depletion (n-o), as quantified by counting the number of cells per cluster (p, n = 14, 13, 15, 14). Statistical tests: Kruskal–Wallis with post-hoc multiple comparison analysis. Data are presented as mean values ± SD.

## Notes

### Competing Interest Statement

The authors have declared no competing interest.

